# Predicting splicing patterns from the transcription factor binding sites in the promoter with deep learning

**DOI:** 10.1101/2023.04.09.536141

**Authors:** Tzu-Chieh Lin, Cheng-Hung Tsai, Cheng-Kai Shiau, Jia-Hsin Huang, Huai-Kuang Tsai

## Abstract

**Background:** Alternative splicing is a crucial mechanism of post-transcriptional modification responsible for the transcriptome plasticity and proteome diversity of a metazoan cell. Although many splicing regulations around the exon/intron regions have been discovered, the relationship between promoter-bound transcription factors and the downstream alternative splicing remains largely unexplored.

**Results:** In this study, we present computational approaches to decipher the regulation relationship connecting the promoter-bound transcription factor binding sites (TFBSs) and the splicing patterns. We curated a fine data set, including DNase I hypersensitive sites sequencing and transcriptome in fifteen human tissues from ENCODE. Specifically, we proposed different representations of TF binding context and splicing patterns to tackle the associations between the promoter and downstream splicing events. Our results demonstrated that the convolutional neural network (CNN) models learned from the TF binding changes in the promoter to predict the splicing pattern changes. Furthermore, through an *in silico* perturbation-based analysis of the CNN models, we identified several TFs that considerably reduced the model performance of splicing prediction.

**Conclusion:** In conclusion, our finding highlights the potential role of promoter-bound TFBSs in influencing the regulation of downstream splicing patterns and provides insights for discovering alternative splicing regulations.

## Background

Gene splicing endows the transcriptional diversity of the metazoan genome. Splicing is the process by which introns are removed from the nascent pre-mRNA and exons are joined, generating the functional mRNA. Alternative splicing (AS), the selective removal of exons and reconnection of exons by multiple processes, is known to play a pivotal role in regulatory pathways from invertebrates to mammals [1, 2]. By the regulatory mechanism of AS, a single gene is capable of generating multiple RNA molecules encoding proteins with different functions [3]. The importance of AS lies in the evidence that the human genome has been estimated more than 95% of multi-exon genes undergo alternative splicing in an underlying tissue-specific manner [4]. Moreover, the variations in splicing patterns are prevalent to associate with many complex diseases in humans [5, 6], and one-third of all disease-associated alleles have been estimated to alter splicing [7].

Studies on AS regulation have mainly focused on the sequence information of spliced exons and flanked introns. Machine learning has unprecedented performance in predicting exon-inclusion/skipping levels in bulk tissues or single cells. Several computational models to derive “splicing codes” that predict splice site selection in a genomic sequence successfully capture patterns around the skipped exon and elucidate complex regulatory mechanisms from genomic and epigenomic features [8–12]. Despite many efforts to characterize the splicing regulatory codes within the splice sites, the extent and effects of transcription machinery at the relatively distant promoter regions in splicing regulation remain unsolved.

In the past decades, AS has been generally accepted to be tightly coupled with RNA polymerase transcription of the nascent pre-mRNA [13, 14]. Two prevailing models have been proposed to explain the coupling between alternative splicing and transcription: the recruitment model [15, 16] and the kinetics model [14]. Notably, the chromatins are mostly not in linear form; the transcription complex on a promoter affects the recruitment of splicing factors and elongation of RNA polymerase II to promote exon exclusion through chromatin looping [17]. In addition, various DNA-binding proteins have been reported to influence the AS patterns by changing epigenetic conditions in the promoter [18].

Each gene contains a set of unique combinations of TF binding sites (TFBSs) in the promoter that determines its temporal and spatial expression. Transcriptional regulation is usually a combinatorial effect of multiple TFs binding to *cis*-regulatory elements located in the proximate and distal regions from transcription start sites [19]. Date to 20 years ago, the regulation of exon splicing patterns was demonstrated directly through the specific TFBS occupancy in the promoter [20, 21]. Moreover, the coupling of promoter and splicing is later proposed with extensive regulator mechanisms [22, 23]. Given the three-dimensional folding of chromatin loops, the proximal promoter- or distal enhancer-bound factors joined into transcription compartments correlate with alternative splicing of exons [24]. Although the biological findings connect the promoter with AS by focusing on a few gene models, the hypothesis that promoter architecture in terms of TFBS composition regulates AS remains unexplored at the genome-wide level.

In this study, we developed analytical strategies to approach this question using data of both RNA-seq and DNase-seq in pairs across the different human tissues from the ENCODE project. We first considered the associations between the occurrences of more than 300 TF binding motifs in the promoter and the corresponding splicing patterns. Secondly, we examined whether the changes in TF binding condition were able to predict the splicing change by studying the relative changes of the splice-in percent (PSI) values between any paired tissues. Then, we conducted machine learning methods and deep learning neural networks to predict the splicing patterns. Notably, the convolutional neural network (CNN) models that took complete TF occupancy information in promoter regions as input achieved the highest performance at 0.889 of the area under receiver operating characteristic curve (AUROC). Lastly, we applied the importance analysis of the CNN models for each TF and identified some important TFs that affecting the splicing prediction genome-widely.

## Results

In this study, we considered the cassette exon splicing, which is the most frequent alternative splicing type in the human genome [36]. We proposed two scenarios to examine the relationship between TFBSs in the promoter and the splicing patterns of the gene. First, we asked if compositions of TFBS occupancies, which were defined as the expressed TFs (TPM > 1) in the given tissues and their binding motifs in the open chromatin regions, are associated with the splicing patterns of the gene. Second, we asked if the changes of TF binding condition in the promoter modify the splicing efficiency of the cassette exon usage by comparing their PSI values. The data preprocessing procedures for TFBS identification in the promoter and exon-skipped events are illustrated in Fig. 1A. The TF binding profiles of each promoter were curated by integration of DNase-seq for open-chromatin regions, human TF motif scan, and expression profile across 15 tissues. The splicing patterns of each gene were analyzed based on the transcriptome in different tissues.

**Figure 1.**
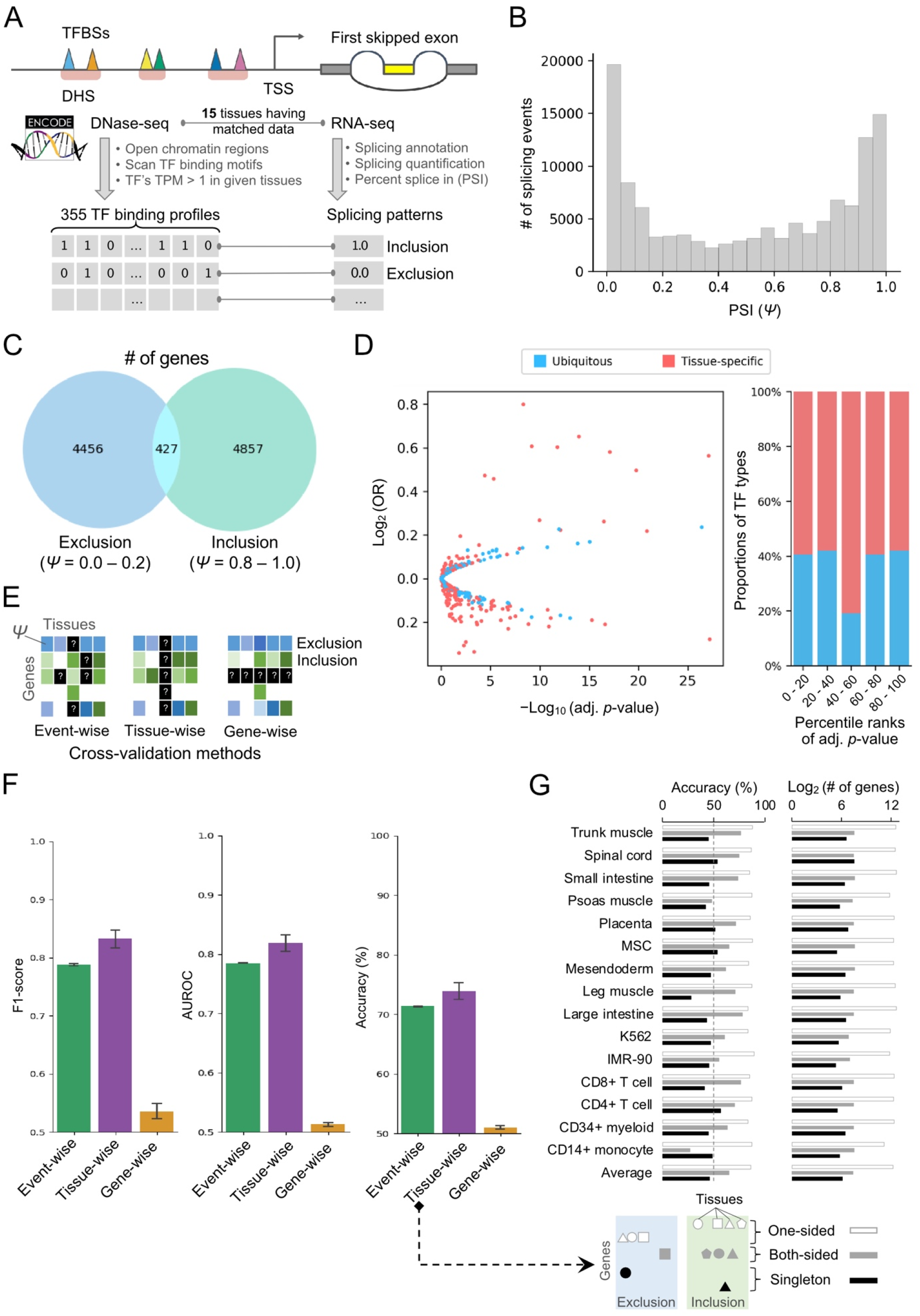
(A) The workflow schema and the experiment design. We obtain 15 tissues that have matched DNase-seq and RNA-seq from ENCODE. DNase-seq data was used to identify open chromatin regions and followed by TF motif scanning to identify TF binding profile in promoter. RNA-seq data was processed by the MISO program to obtain percent splice in (PSI) metrics which represent the splicing pattern of the first skipped exon. (B) PSI distribution histogram. The horizontal axis represents the PSI value and the vertical axis represents the number of skipping exon events. (C) Venn diagram of the exclusion group gene and inclusion group gene. The exclusion group gene defined as PSI < 0.2 and the inclusion group gene defined as PSI > 0.8. (D) Volcano plot of the chi-square test results and the TF expression tissue specificity distribution along with ranking *p*-values of the chi-squared test. The horizontal axis of the volcano plot represents the -log_10_ (adjust *p*-value) and the log_2_ (OR). The chi-square test *p*-value is corrected by Bonferroni multiple test correction. The blue dot denoted the ubiquitously expressed TFs (tau < 0.8) and the red dot denoted the tissue-specific expressed TFs (tau ≥ 0.8). (E) The schema of validation strategies. From left to right represents event-wise, tissue-wise, and gene-wise validation schema, respectively. (F) The model performance of event-wise, tissue-wise, and gene-wise validation schema. For left panel to right panel represents F1-score, AUROC, and accuracy, respectively. (G) The gene were assigned into three groups according to the splicing forms across all tissues. One-sided denotes the genes belonging to same splicing form in more than two tissues; both-sided denotes the genes having both inclusion and exclusion forms in 15 tissues; singleton denotes the genes expressed in a particular tissue only. The accuracies of prediction and number of genes in three groups were calculated respectively for each tissue from the tissue-wise validation experiments.

### Characterizing the TFBS occupancies in the promoter and first cassette exons across tissues

We investigated the associated relationship between the TFBSs in the promoter and the first cassette exon, which is relatively closed to the promoter. The distribution of the PSI values as exon usage levels was bimodal across 15 human tissues (Fig. 1B). Here, we defined the PSI values smaller than 0.2 and larger than 0.8 as the exclusion form and inclusion form, respectively. Based on the criteria, the usage of the first cassette exons of human genes across 15 tissues was mostly skewed in either one of the categories, *i.e.,* exclusion or inclusion forms (Fig. 1C). There were only 4.6% of genes having both splicing forms in different tissues.

Experimental studies have shown that the promoter architecture, by using different gene promoters, affects the splicing patterns of the exon skipping in the gene bodies [37, 38]. Following this idea, we sought to examine whether the promoter architecture in terms of TFBS occupancies as the features determine the inclusion or exclusion of the first cassette exon. First, we asked which TFBSs were predominant within the promoters of these genes with different splicing patterns of their first cassette exon. In order to address this, the discrepancy between the frequency of individual TFBS on the promoters of the exclusion sets and that of the inclusion sets was evaluated independently by using a chi-squared (χ^2^) test for each tissue. Considering an adjusted significance level of *p*-value < 0.001 after Bonferroni correction, more than half of TF binding motifs are significantly enriched in the promoter of either exclusion or inclusion sets. In addition, we calculated the gene expression specificity index tau [31, 32] for each TF and set 0.8 as the cut-off for tissue-specific TFs. However, there is no particular enrichment of TFs showing more enriched across statistical significance ranks (Fig. 1D, right panel).

Next, we considered the complex relationship among TFBSs within promoters on the prediction of splicing patterns by using a machine learning approach. We employed the XGBoost method [39], a decision-tree-based ensemble model, and used the presence of TFBSs within the open chromatins of promoter as input data to predict the inclusion or exclusion of the first cassette exons. Due to the coarser resolution of DNase-seq and *in silico* motif scanning to profile the TFBS occupancies, we noticed that some genes share identical features in different tissues. We thus removed the samples that share identical features in the training data from the testing data of the given tissues to avoid the fallacy of prediction accuracy in the cross-tissue evaluation. Herein, we proposed three different cross-fold validation schemes in order to properly evaluate prediction performance (Fig. 1E). For event-wise scheme, we randomly left 10% of promoter-splicing pairs as the independent testing data and performed a 10-fold cross-validation (CV). For tissue-wise scheme, we conducted leave-one-tissues-out cross-validation by treating the promoter-splicing pairs from a single tissue as the independent testing data. For gene-wise scheme, we used 90% of genes with all promoter-splicing pairs across tissues to train model and remained 10% of genes were for an independent testing set. In Fig. 1F, three evaluation metrics, including F1-score, AUROC, and accuracy, were shown to compare the prediction performance in different CV schemes. Interestingly, the prediction performance using event-wise scheme achieved an F1-score and AUROC closed to 0.80 (Fig. 1F, green bars). In the cross-tissue validation results, we further observed that the overall performance of the models obtained an average AUROC of 0.84 (Fig. 1F, purple bar). However, all three metrics underlying gene-wise CV could yield slightly better than random guess at 0.50 (Fig. 1F, yellow bars).

It is worth noting that the gene-wise CV scenario indeed examined whether the generalization of a trained model enables to classify the splicing events using the unseen promoter information about TF binding profiles, which were not included in the training dataset. We later addressed a following question if the same gene promoter in different tissues both present in the training and testing sets was critical for prediction performance. Subsequently, we split the genes into three groups, i.e., one-sided, both-sided, and singleton, according to their splicing forms across all tissues and re-examined the results of prediction accuracy in the individual tissues. In contrast to the genes with one-sided and both-sided splicing forms, the trained models using data from other tissues did not predict the splicing forms of the singleton genes correctly in the given tissue (Fig. 1G, left panel). Furthermore, we counted the number of genes in the respective groups (Fig. 1G, right panel), and found that a good overall performance of the models underlying tissue-wise CV was dominant by the large number of genes with one-sided splicing form across all tissues. The poor prediction on those small portions of singleton genes (less than 200) did not cause a drastic drop in overall prediction accuracy. In summary, our current approach failed to construct the models with generalization ability to infer the splicing forms using promoter information that pertains to TF binding profiles.

### Changes of TF binding to the promoter reflect the distinct exon splicing phases

In this section, we sought to examine whether changes of individual TF binding to promoter alter the splicing efficiency that was estimated by PSI values. The PSI value summarizes the splicing condition of the constitutive exons that are included in all or part of transcripts from expressed isoforms [40]. As the fact that ranges of PSI values of different genes are varied across 15 tissues, the genes differ from each other in terms of their efficiency of splicing first cassette exon into the expressed isoforms. As a result, the efficiency of exon usage should be considered for each gene itself instead of the absolute PSI (Ψ) value. To this end, we applied the Z-score transformation to normalize the absolute PSI scores of all genes. Of note, some genes that had a smaller PSI range (< 0.2) and/or expressed in less than three tissues were discarded in the following experiments. We then defined the top 20% and last 20% of transformed *Z*_Ψ_ scores in each gene as the two distinct phases of exon usage, i.e., low and high splicing efficiency respectively (Fig. 2A). To test the hypothesis that changes of TFBS in the open chromatin of the promoter are associated with splicing phase change, the differences of two Z_Ψ_ and their TF binding occupancies in a given paired tissues for each gene were calculated (Fig. 2B). The distribution of delta Z_Ψ_ scores was shown in Fig. 2C, where the unchanged group (same splicing phase) was below 1 and the changed group (different splicing phase) was larger than 1.8. Of note, no overlapped events were observed between concordance and discordance groups.

**Figure 2.**
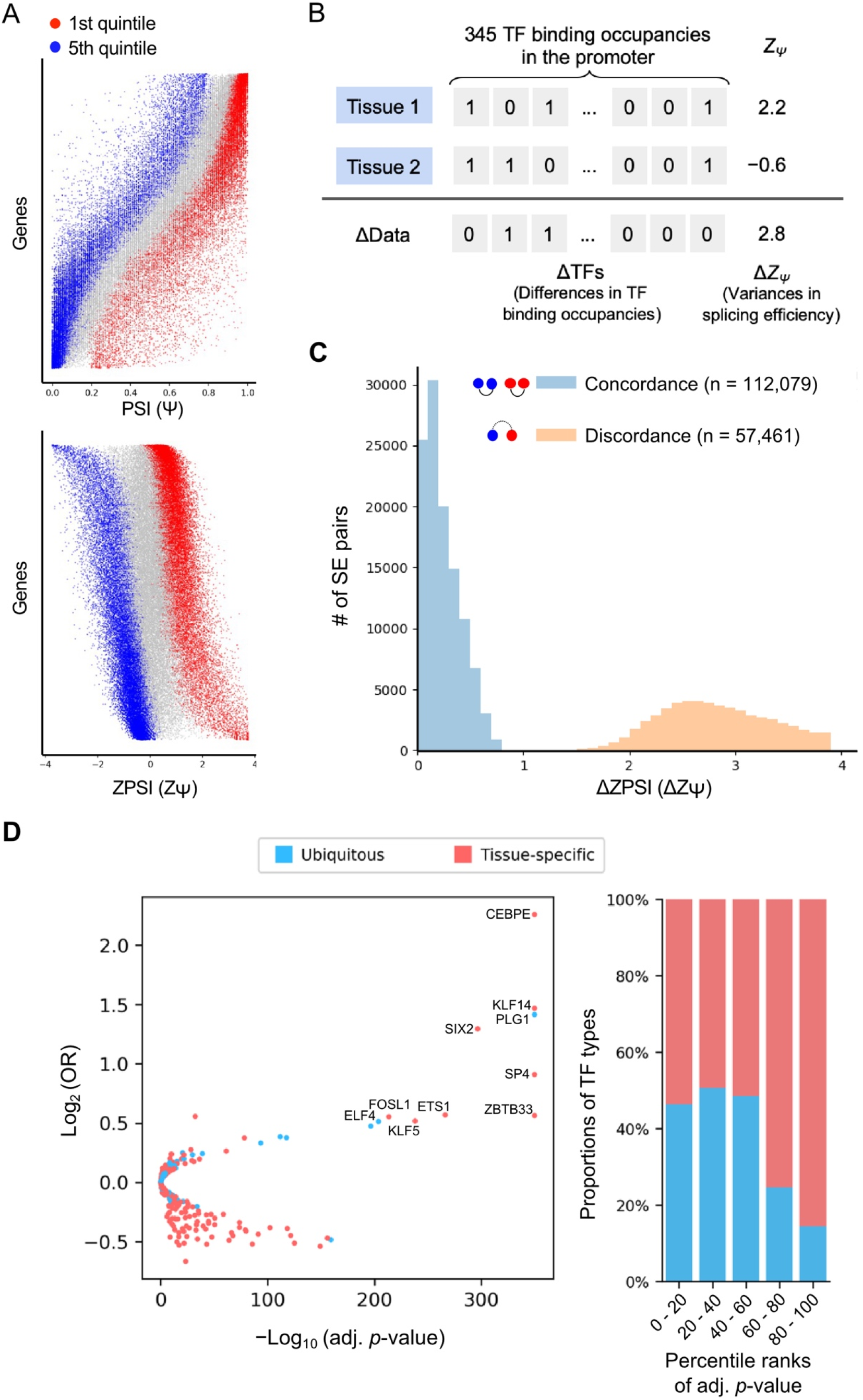
(A) The distribution of PSI and ZPSI. Each horizontal line represents PSI values of a gene and the vertical axis was sorted by gene median PSI. The blue dots denote the first quintile (top 20%) of PSI and the red dots denote the fifth quintile (latest 20%) of PSI. (B) The “delta” schema of splicing events. For each gene, we enumerate all tissue pairs and perform exclusive-or (XOR) operation on the TF binding occupancies and yield ΔData representation which means the differences in TF binding occupancies. For the splicing pattern, we calculate the absolute difference of the ZPSI and yield ΔZ_Ψ_, which represents the variances in splicing efficiency. (C) The distribution of ΔZ_Ψ_ among splicing status unchanged group (concordance) and changed group (discordance). The distribution showed a clear bimodal pattern, that the discordance ΔZ_Ψ_ is distinctly higher than the concordance ΔZ_Ψ_. (D) The chi-squared test of association between TFBS-occupied differences and splicing phases. The left panel is the volcano plot of the chi-square test; the horizontal axis represents the -log_10_ (adjusted *p*-value) and the vertical axis represents the log_2_ (OR). Top 10 significant TFs are shown in their names. The right panel is the ratio of tissue-specific and ubiquitous TFs among adjusted *p*-value rankings.

To examine the association between TFBS-occupied difference and splicing phase for individual TFs, we constructed a 2 × 2 contingency table for each TF. Specifically, for each tissue pair in one gene, we assigned the pair into groups according to whether its TFBS occupancy is changed (ΔTF*α* = 0 or ΔTF*α* = 1), and whether the splicing phase is changed (concordance or discordance). We thus calculated the odds ratio from contingency table and applied chi-squared test. About two-third of TFs, their binding occupancy changes were significantly associated with splicing phase changes (N = 203, adj. *p*-value < 10^-3^, Fig. 2D). Since every tissue usually expresses different sets of TFs to control the cell fate [41, 42], we estimated the tissue specificity of TF expression by tau score [32]. More than half (53%) of TFs among those non-significant groups were ubiquitously expressed, while most of the TFs (75%) among those significant associations with splicing phase change were tissue-specifically expressed (Fig. 2D). Of note, the open chromatin regions in the promoter of the same gene in different tissues show less variations. Thus, TFBSs without filtered by expression profiles of given TFs did not show any significant association. Therefore, although the DNA sequences of the promoter are identical, the divergence on the TF expression across different tissues is a likely regulating mechanism to affect the splicing phase change.

### Machine learning confirm the association between TF binding changes and splicing phase shift

Next, we employed different machine learning algorithms, including logistic regression, XGBoost (ensemble tree algorithm), and deep neural network methods, to test whether the combinations of TF binding changes predict the splicing phase changes. To monitor sensitivity and specificity simultaneously, we assessed the models using the AUROC in the plot of the true positive rate (TPR) against the false positive rate (FPR) for five-fold cross-validation tests (Fig. 3A). Three classifiers achieved an average AUROC of 0.691, 0.766, 0.771 for logistic regression (LReg), deep neural network (DNN), and XGBoost (XGB) models, respectively on all the events of the dataset. Since there were imbalanced data sets in the changed and unchanged groups, the area under the precision-recall curve (AUPRC) is also instructive to assess the model performance (Fig. 3B). The XGB models also achieved a greater mean AUPRC of 0.630 than 0.531 and 0.624 respectively for LReg and DNN. Because there is often more than one binding site in the promoter for each TF, we also constructed other ML models using frequencies of all possible TF binding site changes between promoters as the features. The overall performance of prediction of splicing phase change was decreased about 6% based on AUROC. This indicates that the decision tree-based ML method could not deal with the frequencies of TFBSs change properly.

**Figure 3.**
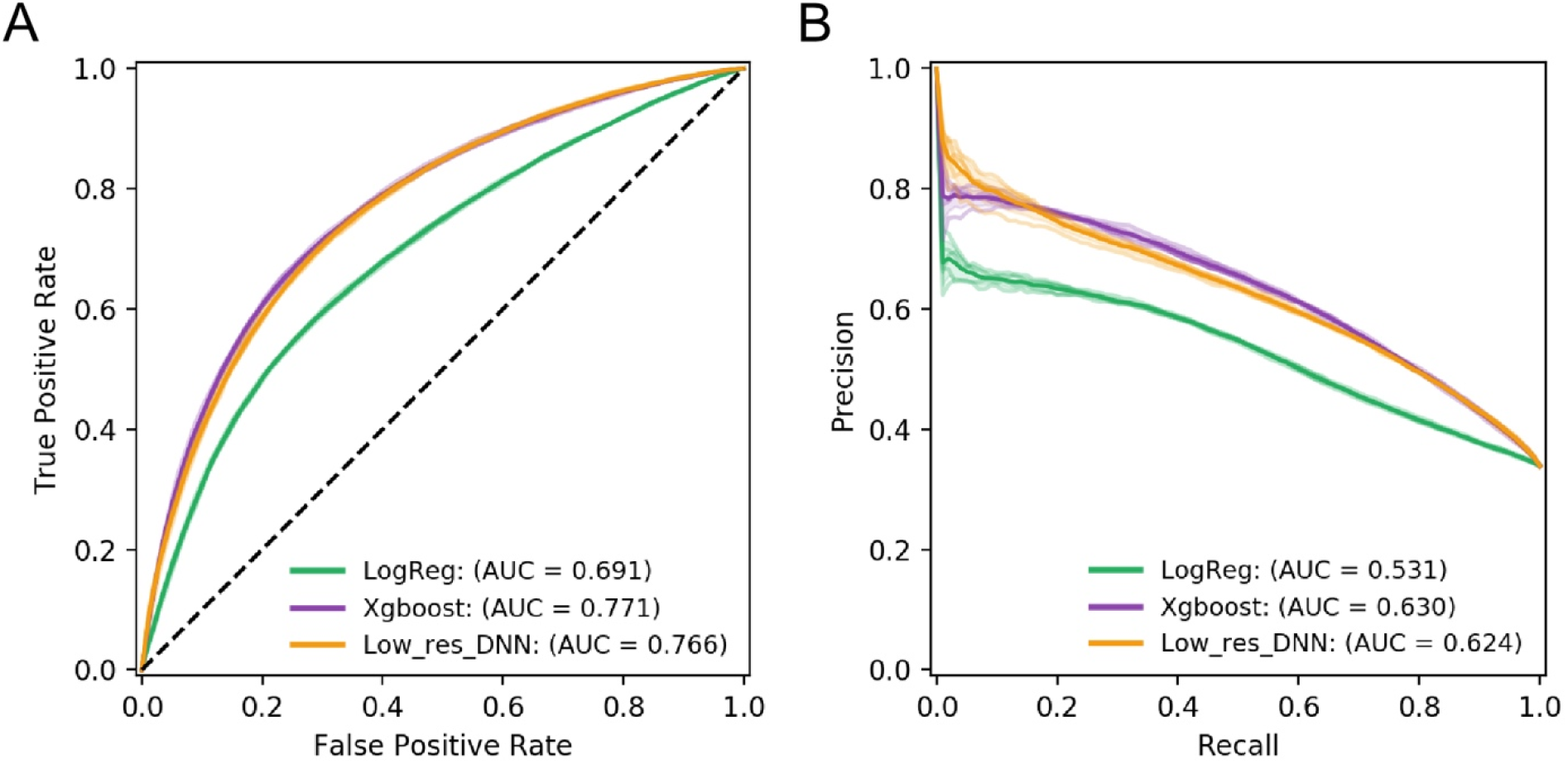
(A) The area under receiver operating characteristic curve (AUROC) of Logistic regression, XGBoost, and low-resolution deep learning model. The input of the low-resolution deep learning model only contains a single array of TF occupancy information denote as low-resolution. Of note, the XGBoost model has the highest AUROC. (B) The area under precision-recall curve (AUPRC) of Logistic regression, XGBoost, and low-resolution deep learning model. With the same trend of AUROC, the XGBoost model has the highest AUPRC.

### Integration of TFBS locations in the promoter using deep learning models improve prediction performance

We next integrated the position information of TFBSs in the promoter as the features to train the deep neural network (DNN) and convolution neural network (CNN) models respectively. The two-dimension array consisting of 2,500 bp and 345 TF binding changes were used as the input features as shown in Fig. 4A. The architecture of the CNN model includes the one-dimensional convolutions kernels, which are designed as the filters for revealing the combinations of TF binding changes. The convolution layers are followed by a max-pooling layer with sliding window size and a stride step of 10 units. And a single flatten layer with 256 neurons was used to summarize all features and followed by three hidden layers. To prevent overfitting, the dropout technique was applied to remove 25% of the connected neurons in the flatten and hidden layer during the training (26).

**Figure 4.**
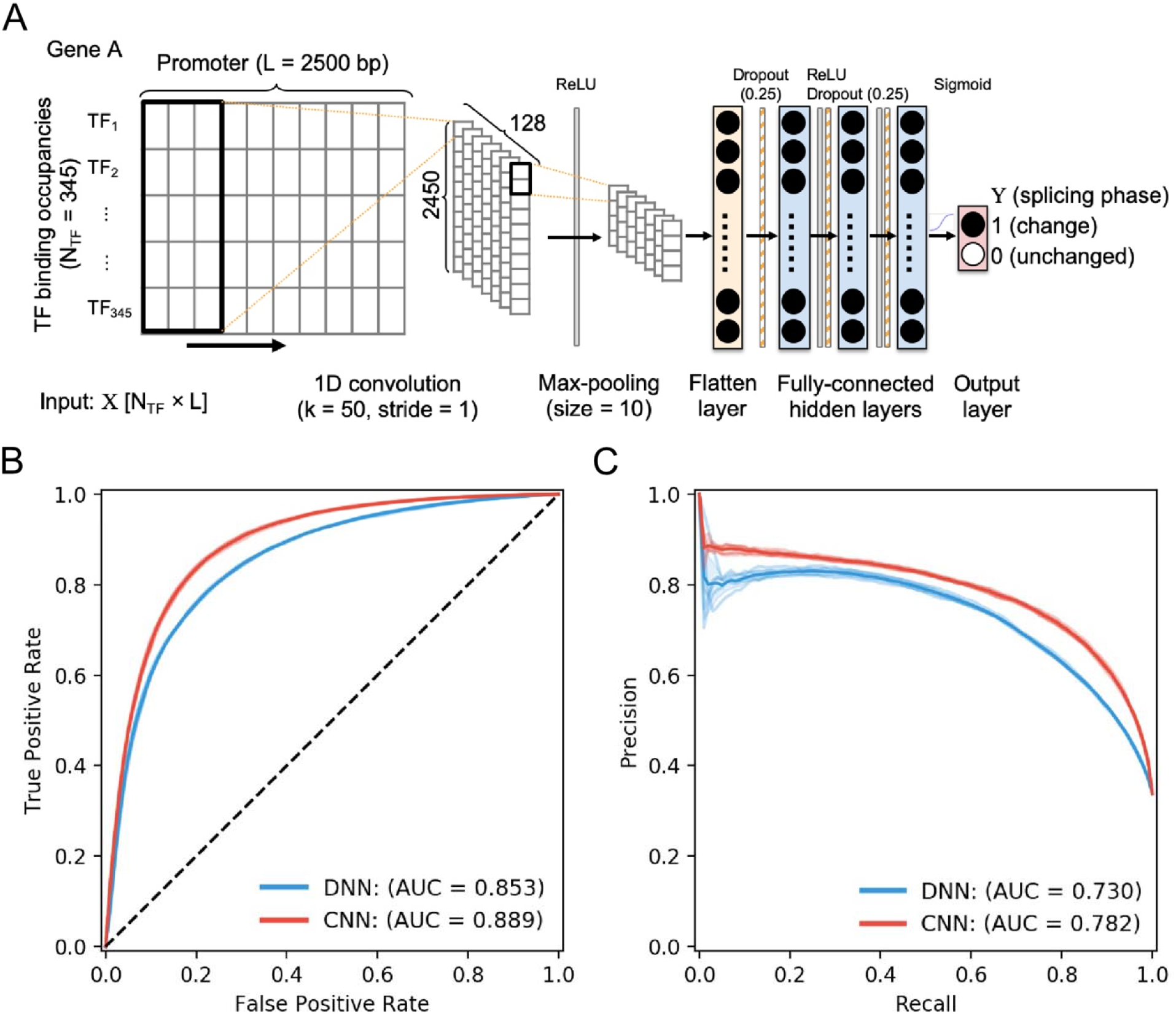
(A) The convolutional neural network schema. The first layer is a convolution layer with ReLU activation function and followed by a max-pooling layer. After pooling a flatten layer was applied to reshape the input. Then three dense layer is added followed by a sigmoid function to classified the output. (B) The area under receiver operating characteristic curve (AUROC) of convolutional neural network (CNN) and deep neural network (DNN). (C) The area under precision-recall curve (AUPR) of CNN and DNN. Both AUROC and AUPR suggest the CNN has the better performance.

Training the network with input matrices including both TFBS and their interactions with other TFs markedly impacts the performance of the splice predictions. In contrast to the performance of previous DNN models using only TFBS changes input (Fig. 3A), current DNN classifiers achieved greater AUROC, increasing from the average 0.766 to 0.853 (Fig. 4B). The CNN classifiers achieved an even greater AUROC of 0.889 (Fig. 4B). Additionally, CNN models achieved greater AUPRC for all five-fold experiments than DNN models, increasing the average from 0.730 to 0.782 (Fig. 4C).

### Evaluation of TF changes on the splicing patterns

We next to understand the importance of TF motifs on splicing patterns utilized by the network to achieve its remarkable accuracy. In brief, we performed systemic *in silico* substitution of each TF change as zero, then measured the effects on the CNN model’s prediction. The importance of each TF was estimated by the fraction of changed prediction under the *in silico* substitution. The underlying idea is if assume a TF plays a key role in regulating splicing patterns, the prediction output of the machine learning model should change dramatically after substitution rather than other TF. We performed importance analysis on each TF and ranked them by their importance measurement, and found that a small proportion of TFs resulted in dramatical changes in the splicing prediction (Fig. 5A). As most of TFs had a little effect on the CNN model performance, we highlighted top-ranked 19 TFs with outlier values based on the interquartile range rule (Q3 + 1.5 × IQR) as the candidate splicing regulators.

**Figure 5.**
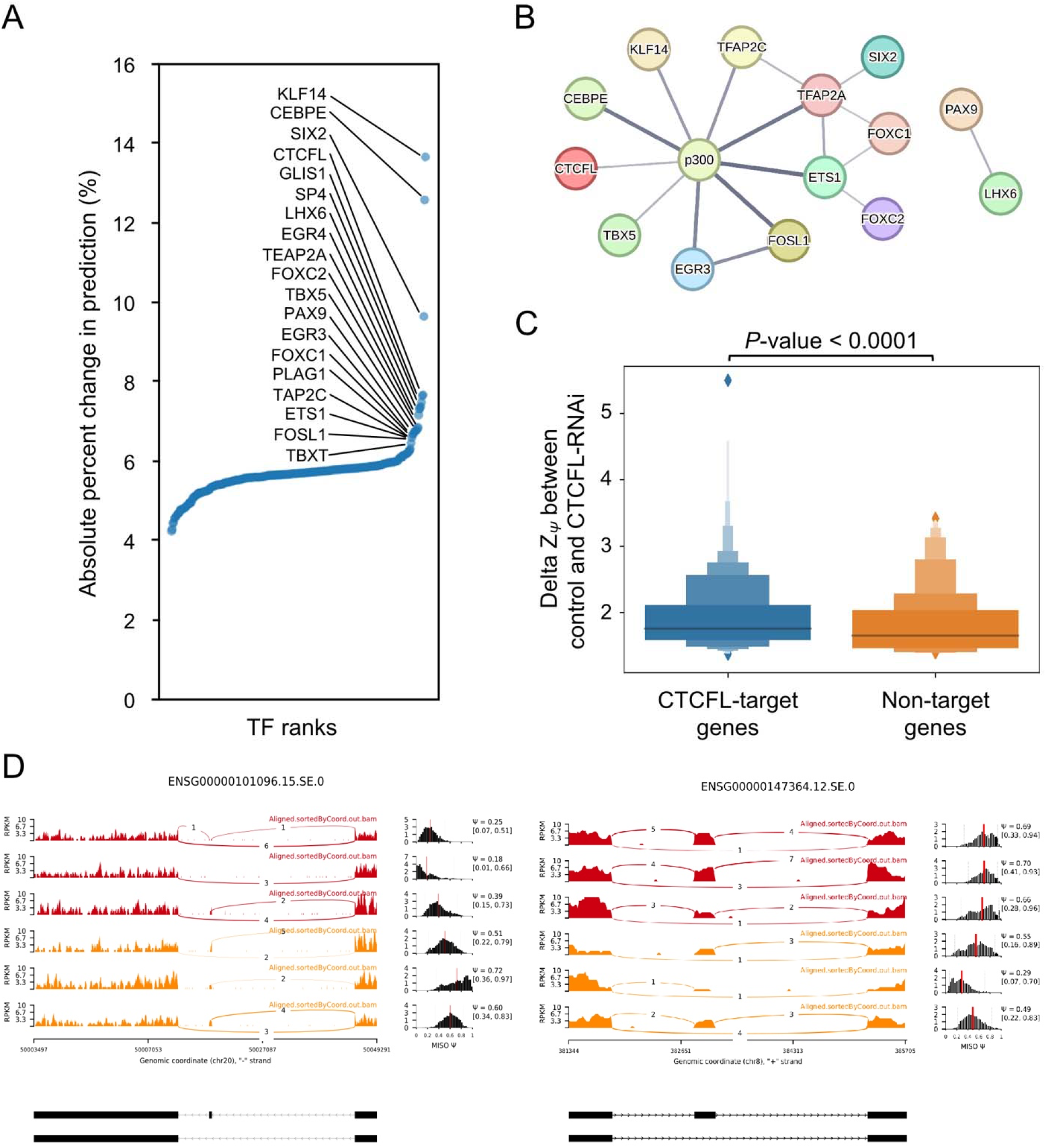
(A) The rank order plot of importance analysis. The horizontal axis represents the TF importance ranks. The vertical axis represents the importance measures (see importance analysis in method section). (B) The gene association network was constructed from the STRING database for top important TFs with p300. The thickness of edges denotes the strength of data support according to textmining, experiments, and databases. (C) The distribution of ΔZ_Ψ_ between control and CTCFL-RNAi experiment. The ΔZ_Ψ_ values of the CTCFL-target genes show significant differences than that of non-CTCFL target genes with Wilcoxon rank-sum test (*p*-value < 0.0001). (D) The sashimi plot and PSI distribution across control and CTCFL-RNAi experiment. The left panel shows the first skipped exon event of ENSG00000101096. The right panel shows the first skipped exon event of ENSG00000147364. Red samples were from the control of CTCFL experiments and orange samples were from the CTCFL-RNAi samples.

Previous studies have demonstrated that binding of the acetyltransferase p300 at promoter regions modifies acetylation of splicing factors, and thereby modulate the alternative splicing pattern of the gene [43, 44]. We thus submitted our 19 candidate TFs and p300 to the STRING database [45] for identification of their interactions. We applied default settings to search both functional and physical protein associations with medium confidence score of 0.400 in the STRING database (ver. 11.5). Then, we configured the network between query proteins only to reveal the associations among them. Interestingly, the network was relatively less complex and p300 were thought of as a hub gene associated with nine out of 19 top-ranked TFs (Fig. 5B). Moreover, the interaction between KLF14 and p300 is experimentally and functionally confirmed that the binding of KLF14 to the promoter recruits p300 to increase the levels of acetylation associated with transcriptional activation [46]. Although the interaction between KLF14 and p300 on the gene activation was not investigated in the context of splicing, compelling evidence showing a direct link between histone modification and splicing [17, 18] raises the intriguing possibility of KLF14/p300 complex in modulating exon splicing. Similarly, some top-ranked TFs might share a common mechanism in regulating RNA splicing via recruitment of p300 to promote the deposition of histone acetylation at the promoter.

Lastly, to further confirm our *in silico* prediction for potential splicing regulators, we obtained the K562 CTCFL shRNA knock-down RNA-Seq data [47] and its control from previous research [48]. We re-analyzed the splicing status by calculating PSI through MISO and applied Z-score transformation using the previous method in machine learning model training. We observed the ΔZ_Ψ_ values of CTCFL-target genes were higher than that of non-target genes significantly (Fig. 5B, with *p*-value < 0.0001, Wilcoxon rank-sum test). This revealed in the CTCFL deplete condition, genes targeted by CTCFL change their first skipped exon usage thus influence ΔZ_Ψ_. We further seek for case studies to investigate how splicing status changed in CTCFL-target genes under CTCFL depletes (Fig. 5C). The first skipped exon in ENSG00000101096 has a higher skipped exon usage and increases the average PSI value. In contrast, in ENSG00000147364 the first skipped exon usage reduced in the CTCFL deplete condition thus has a lower average PSI value. These results suggest that CTCFL can influence the splicing pattern. Nevertheless, CTCFL shows a dual function in splicing regulation, not only increase skipped exon usage but also reduce usage in some genes. This result also matches the previous study on CTCFL-depletion mediate alternative splicing change in MCF7 cell line [49]. In the CTCFL-depletion they detect exclusion of 361 and the inclusion of 221 alternative exons compared to the normal condition. The CTCFL can influence the recruitment of RNAPII and thus impact the RNAPII elongation speed and finally alter the splicing result of pre-mRNA. Overall, these results support the feasibility of our modeling and importance analysis approaches for *in silico* prediction.

## Discussion

The applications of machine learning methods to characterize the regulatory potential of genomic sequences on alternative splicing have been a subject of interest for over a decade [8, 50]. Instead of using the genomic information around the splicing exons, in this study, we focused on the upstream promoter region for predicting downstream exon-skipped events genome-widely. In contrast to some previous study using the DNA sequences directly [8, 9, 11], one major difference of our approach is that we applied TF binding motif scan with prior domain knowledge to represent the sequence information in the promoter. We demonstrate how the promoter signals in terms of TFBS profiles can be integrated using machine learning approaches for the further implication of association between the promoter and alternative splicing. Our results showed that the prediction accuracy differed among the different algorithms and input information. Notably, one-dimensional CNN architecture is highly capable of learning the regulatory code from the TF binding changes in the promoter to discriminate the splicing patterns (Fig 4).

The main drawback of this study is the limited number of tissues because we aimed to use a high-quality dataset to avoid the noise and artifacts in the DNase-seq and RNA-seq datasets conducted by different labs. Thus, we excluded any experiments that did not meet every quality standard defined by ENCODE. When conducting the data analyses, we noticed that the splicing forms for most of the gene were not varied extensively in these 15 tissues (Fig. 1C). Inspired by the previous study to avoid fallacy of model performance using alternative cross-fold validation schemes properly [51], we implemented three different CV schemes, i.e., event-wise, tissue-wide, and gene-wise, to evaluate generation performance carefully. In the course of examining the difference across three CV schemes to find possible reasons for high performance in the tissue-wise evaluation, we noticed that majority of genes were expressed in more than two tissues and displayed same splicing form. Because every gene promoter in different tissues shares most TFBS features, the event- and tissue-wise schemes are subject to the problem of test set contamination and could lead to an artificially inflated accuracy in this study. On the bright side, there is considerable room for improvement in model generalization by collecting varied splicing forms of every gene from different tissues extensively to evaluate promoter-splicing interactions.

To address the problem of shared TFBSs in promoter across tissues, we turned to look at the TF binding changes in promoter (Fig 2B). Notably, this approach diminished the high similarity of TFBS features in tissues and making a comparison in any given paired tissues also augmented the datasets incrementally for improvement of the model training. On the other hand, we considered the changes in splicing efficiency (ΔZ_Ψ_) by introducing a transformation procedure of absolute PSI values into the efficiency of exon usage. Our computational method is different than a previous study using the absolute PSI values to estimate splicing efficiency directly [52]. The fact that the ranges of the PSI values in a particular gene across 15 tissues are mostly ununiformed distribution is evident as the averaged PSI values of genes from closed to 0 or 1 (Fig 2A). The Z-transform method could remain commensurate in the scale to measure splicing efficiency for each gene accordingly. In addition, instead of using fixed arbitrary cutoff values (e.g., Ψ < 0.2 and Ψ > 0.8) to subsect the splicing status, we applied a percentile threshold to divide genes into two tendencies, i.e., “splice-in” or “splice-out”. This approach avoids that those small-PSI-ranged genes are skew to be classified into a single group of splice-in or splice-out. Based on our observation, it is perhaps noteworthy to rethink about the definition of the splicing status using PSI as a metric to explore alternative solutions in discovery of splicing mechanisms. By carefully considering the fundamental issues in our preprocessing procedures on data, this study provides a different perspective to study how TFs in promoter affects the exon splicing genome-widely.

To train the prediction model of splicing phase shift, we used two different input data, *i.e.*, an array of TF binding changes and a matrix of full TF binding changes along with the promoter regions. Our results demonstrated that training the DNN models with varying input of TF binding context noticeably impacts the accuracy of the splicing phase shift prediction (Fig. 3 and 4). Despite amount of trainable network parameters drastically are increased when using an input of TF binding context, DNN models is capable to automatically learn the task from the training data. Remarkably, CNNs achieved even higher prediction performance than DNNs with matrices of TF binding context (Fig. 4). In contrast to DNNs, CNNs indeed are designed to deal with high-dimensional inputs by applying of a serious of convolutional and pooling steps [53, 54]. A likely explanation for high accuracy boosting in CNNs is the convolutional operations, which learned higher-level features from the combinations of different TF changes. With the good prediction performance of CNN models, the importance analysis experiments allowed us to identify a couple of TFs that potentially involve in splicing regulation. To our knowledge, our study is the first genome-wide effort to investigate that the splicing pattern changes across tissues were accurately predicted from the TF binding occupancies in the promoter.

## Materials and Methods

### Data processing and sample selection

We downloaded both the DNase-seq peak BED files and the RNA-seq data for 15 human tissues from the ENCODE data portal [25]. To obtain high quality of data, the data without any flags, such as insufficient read depth, in the experimental metadata that were reported by the ENCODE Data Coordination Center are used in the following experiments. For DNase-seq datasets, the standard pipeline (accession: ENCPL201DNS for single-ended data, ENCPL202DNS for paired-ended data) from ENCODE called the peaks using hotspot2 algorithm with 1% false-discovery rate. For RNA-seq data, the ENCODE RNA-seq pipeline for long RNAs (accession: ENCPL002LSE for single-ended data, ENCPL002LPE for paired-ended data) used the STAR program for mapping the reads and the RSEM algorithm for quantification of genes. We used genomic and annotation files of the human reference genome version GRCh37 as provided by release V19 of GENCODE [26].

### Identification of putative *in vivo* TF binding sites

The DNase-seq peaks were used to define the open chromatin regions in the promoter regions (−2 kb to +500 bp from the transcription start site). We downloaded TF motifs from the JASPAR database (ver. 2018) [27] and excluded the fusion TF (i.e., EWSR1/FLI1 fusion) and older versions of motifs from the same TF, as a result, we obtained 407 TF binding motifs from JASPAR. Later, we scanned the sequence from each open chromatin region for each TF binding motif in position-weight-matrix (PWM) format, using FIMO from the MEME (Motif-based sequence analysis tools) suite [28]. Of note, we applied the FIMO with a threshold false discovery rate of < 10^−3^, which is less stringent than the general recommended parameter (< 10^−4^) for putative *cis*-regulatory elements detection. Since we only considered TF binding sites located in the open chromatin regions, the general parameter is too stringent for our purpose.

### RNA-seq processing and calculation of cassette exon usage (PSI)

To estimate the splicing level for each exon and tissue, we first used CATANA [29] to annotate AS events in all human transcripts for the AS annotation index file. The BAM files of RNA-seq data generated by the ENCODE were used to estimate the percent spliced-in (PSI) values for the first cassette exon of the protein-coding genes using the MISO (Mixture of Isoforms) tool [30]. For the calculation of the *Z_Ψ_* score, we first selected the genes that PSI range is larger than 0.2 across different tissues and then standardized their PSI by z-score transformation for each gene.

### Enrichment analysis

We analyzed the association of TF binding occupancies and splicing patterns from 2 × 2 contingency tables categorizing all human genes according to the occurrences of binding sites for a given TF and splicing patterns (exclusion or inclusion in Fig. 1D). In parallel, we built the contingency table to analyze the association between TFBS-occupied differences and splicing phases (concordance or discordance in Fig. 2D) for each TF. The odds ratio (OR) based on the contingency table was calculated for each TF and a chi-squared (χ2) test was applied to determine the statistical significance of the association. The *p*-value is adjusted by Bonferroni correction (and its −log10 transformation) for the association, and the odds ratio with log2 transformation is a measure of the effect size. The adjusted *p*-value < 0.001 is considered as significant.

### Tau index of TF tissue specificity

We calculated the tissue specificity index tau [31, 32] using the gene expression of each TFs across different tissues, as follows:

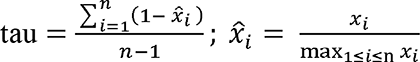

where *x_i_* represents the gene expression of TF *x* in tissue *i*; and *n* is the number of tissues expressing the TF (TPM > 1). We then adopted the cut-off of tau based on a previous study [33] and defined the TFs with tau ≥ 0.8 as tissue-specifically expressed.

### Machine learning and deep learning models

In order to get a better prediction power, we compared the accuracy between four methods, logistic regression, XGBoost, deep neural network (DNN), and convolutional neural network (CNN). To avoid biases caused by imbalanced data, we applied a balanced sampler as the concept described on the imbalanced-dataset-sampler (from https://github.com/ufoym/imbalanced-dataset-sampler) to our training dataset before model training. We trained the basic logistic regression model with default parameter settings described in the scikit-learn [34]. For the XGBoost model, we limited the max tree depth to 6, set the eta by 1, and used gbtree as a booster.

In this research, we implemented our DNN and CNN models using the PyTorch framework [35]. The architecture of DNN began with flattening the input data and followed by 3 dense layers, with 512, 256, and 128 nodes, respectively. ReLU activation function was applied on the output of each dense layer and then followed by a dropout layer to randomly set 25 percent of input units to 0. The sigmoid function was applied to the final output of the tensor to generate binary classification predictions.

The architecture of CNN is similar to DNN with some modifications. The input data was first processed through a convolution layer which followed by the ReLU activation function, max pooling layer and a dropout layer, and then connected to 2 dense-ReLU-dropout units as described above, both with 128 nodes. The sigmoid function is also used to do the binary classification task.

### Importance analysis

To extract informative TF binding features from the CNN model, we performed an *in silico* perturbation-based analysis to observe the impact on the perturbed input data. Similar to the previous method, we perturb the input by assigning a zero value for a given TF of the input feature (zero-out operation) and perform inference on the trained model. The feature importance through zero-out operation was measured by the output changing ratio. Output changing ratio was defined as N_changed_ / N_total_, where N_changed_ represents the count of changed output label after zero-out and N_total_ represents the total input delta instances number with corresponding TF binding site.

## Availability of data and materials

The source codes supporting the conclusions of this study are available at GitHub repository (https://github.com/bio-it-station/DoTA).

## Authors’ information

### Affiliations

#### Institute of Information Science, Academia Sinica, Taipei, Taiwan

Tzu-Chieh Lin, Cheng-Hung Tsai, Cheng-Kai Shiau, Jia-Hsin Huang, Huai-Kuang Tsai

#### Taiwan AI Labs & Foundation, Taipei, Taiwan

Jia-Hsin Huang, Huai-Kuang Tsai

#### Corresponding authors

Correspondence to Jia-Hsin Huang or Huai-Kuang Tsai.

#### Authors’ contributions

TCL and CHT conceived the study and carried out the bioinformatics pipelines. TCL prepared the initial draft of the manuscript. CKS assisted the data curation. JHH and HKT conceived and designed the research, interpreted the results, and drafted the manuscript. All authors read and approved the final manuscript.

## Ethics declarations

### Ethics approval and consent to participate

Not applicable.

### Consent for publication

Not applicable.

